# Empirical field mapping for gradient nonlinearity correction of multi-site diffusion weighted MRI

**DOI:** 10.1101/2020.05.18.102558

**Authors:** Colin B. Hansen, Baxter P. Rogers, Kurt G. Schilling, Vishwesh Nath, Justin A. Blaber, Okan Irfanoglu, Alan Barnett, Carlo Pierpaoli, Adam W. Anderson, Bennett A. Landman

**Affiliations:** Computer Science, Vanderbilt University, Nashville, TN, USA; Department of Radiology and Radiological Sciences, Vanderbilt University Medical Center, Nashville, TN USA; Department of Biomedical Engineering, Vanderbilt University, Nashville, TN USA; Electrical Engineering, Vanderbilt University, Nashville, TN, USA; National Institute of Biomedical Imaging and Bioengineering, Bethesda MD USA

**Keywords:** Gradient Nonlinearity, Field Estimation, Pre-processing, DW-MRI

## Abstract

**Background:** Achieving inter-site / inter-scanner reproducibility of diffusion weighted magnetic resonance imaging (DW-MRI) metrics has been challenging given differences in acquisition protocols, analysis models, and hardware factors.

**Purpose:** Magnetic field gradients impart scanner-dependent spatial variations in the applied diffusion weighting that can be corrected if the gradient nonlinearities are known. However, retrieving manufacturer nonlinearity specifications is not well supported and may introduce errors in interpretation of units or coordinate systems. We propose an empirical approach to mapping the gradient nonlinearities with sequences that are supported across the major scanner vendors.

**Study Type:** Prospective observational study

**Subjects:** A spherical isotropic diffusion phantom, and a single human control volunteer

**Field Strength/Sequence:** 3T (two scanners). Stejskal-Tanner spin echo sequence with b-values of 1000, 2000 s/mm^2^ with 12, 32, and 384 diffusion gradient directions per shell.

**Assessment:** We compare the proposed correction with the prior approach using manufacturer specifications against typical diffusion pre-processing pipelines (i.e., ignoring spatial gradient nonlinearities). In phantom data, we evaluate metrics against the ground truth. In human and phantom data, we evaluate reproducibility across scans, sessions, and hardware.

**Statistical Tests:** Wilcoxon rank-sum test between uncorrected and corrected data.

**Results:** In phantom data, our correction method reduces variation in mean diffusivity across sessions over uncorrected data (p<0.05). In human data, we show that this method can also reduce variation in mean diffusivity across scanners (p<0.05).

**Conclusion:** Our method is relatively simple, fast, and can be applied retroactively. We advocate incorporating voxel-specific b-value and b-vector maps should be incorporated in DW-MRI harmonization preprocessing pipelines to improve quantitative accuracy of measured diffusion parameters.

## INTRODUCTION

Physics underlying magnetic resonance imaging (MRI) gradient coil designs result in nonuniform magnetic field gradients during acquisition. This leads to spatial image warping [1–4] in magnetic resonance images and gradient distortion in diffusion weighted magnetic resonance imaging (DW-MRI) [5–9]. The introduced spatial variation can impact estimated diffusion tensor information [10] or high-angular resolution diffusion measurements [11]. Bammer et al. show in extreme cases the gradient nonuniformity can lead to an overestimation in the diffusion coefficient up to 30% and an underestimation up to 15% [12]. The severity of the effect increases with distance from the magnet’s isocenter [12] and with higher gradient amplitudes [12, 13]. The artifact becomes especially troubling for multi-site studies that have varying scanner models and manufacturers [14] and for studies utilizing very large gradient amplitudes such as in the human connectome project (HCP) which utilized amplitudes up to 300 mT/m [13, 15, 16]. Recent work has shown the effect of gradient nonlinearities in the HCP cohort results in considerable bias in tractography results and potentially incorrect interpretations in group-wise studies [17].

Various estimates of the coil magnetic field nonlinearities have been applied to improve accuracy within and across sites [18–21]. An adaptive correction of diffusion information proposed by Bammer et al. relies on calculating the spatially varying gradient coil *L*. This approach is achieved by relating the actual gradients with the desired gradients [12], and has become standard practice [22, 23]. However, this approach assumes that the gradient calibration specified by the manufacturer is readily available. Spherical harmonics (SH) based techniques are already implemented by manufacturers in the scanning systems to account for the spatial image warping effects of gradient nonlinearities [1, 24–27]. Yet, the spherical harmonic coefficients are not usually provided to regular users and may be subject to non-disclosure criteria. Additionally gradient nonlinearity correction has been approached using noncartesian MR image reconstruction [28].

To remove the need for the manufacturer supplied specifications, we demonstrate an empirical field-mapping procedure which can be universally applied across platforms as defined by Rogers et al. [29, 30]. At two scanners (scanner A and scanner B), a large oil-filled phantom is used to measure the magnetic field produced by each gradient coil. To estimate the achieved diffusion gradient directions and b-values on a voxel-wise basis, solid harmonic basis functions are fit to the measured magnetic field. The measured diffusivity (MD) and fractional anisotropy (FA) are compared without nonlinearity correction, with nonlinearity correction using estimated fields, and with nonlinearity correction using fields specified by the manufacturer for an ice-water diffusion phantom. The reproducibility is compared between without nonlinearity correction and with nonlinearity correction with the estimated fields for a subject scanned at two positions within the scanner at scanner A. We show that our method removes the need for manufacturer specified spherical harmonic coefficients and that the method reduces MD reproducibility error in-vivo when the effect of gradient nonlinearities is present.

## METHODS

### Measurement of gradient coil-generated magnetic fields

Data were acquired across two 3T scanners: Scanner A and scanner B. Both of these are 94 cm bore Philips Intera Achieva MR whole-body systems and have a gradient strength of 80 mT/m, a 200 T/m/s slew-rate. A phantom is used to estimate the gradient coil fields. The phantom is 24 liters of a synthetic white oil (SpectraSyn 4 polyalphaolefin, ExxonMobil) in a polypropylene carboy with an approximate diameter of 290mm and height of 500mm [30]. This oil is used by the manufacturer for some of their calibration phantoms which made it a reasonable choice. The phantom was placed approximately at scanner isocenter and imaged with a dual echo GRE-based field mapping sequence. Images are acquired at two echo times 1ms apart, and the fieldmap is computed from the phase difference of the two images. This follows the manufacturer’s field mapping and provides a field map with minimal phase wrapping or distortion. Four field maps were acquired, one with shim field set to 0.05 mT/m on each axis X, Y, Z plus a final image with gradient coil shim fields set to zero. Each used a 384 mm field of view with 4 mm isotropic voxel size. Total scan time was approximately 5 minutes. Gradient coil fields were estimated by subtracting the zero-shim field map from each coil’s respective 0.05 mT/m field map. It should be noted that the proposed method requires that the field maps are made using the same coils used to produce the diffusion gradients, and systems that utilize gradient coil inserts may not be able to directly utilize the technique. Field maps were acquired on 40 dates over the course of a year at scanner B while scanner A only one session was acquired with the fieldmapping phantom.

For each coil, we modeled the magnetic field spatial variation as a sum of solid harmonics [12, 31, 32] to 7th order, excluding even order terms due to the coils’ physical symmetry. These basis functions were fit to the field measurements with robust least squares, using all voxels within a 270 mm diameter sphere at isocenter. For comparison, the general shape of the human head is an ellipsoid with an average height of 180 to 200mm [33]. The result was an analytically differentiable estimate of the true magnetic field produced by each gradient coil (Figure 1). This fitting procedure was performed on an average field map derived from a series of scans to ensure stability. On Scanner B, the fitting procedure is also performed on the scanner manufacturer’s estimate of the coil fields as measured during manufacturing and installation. These are provided as a set of solid harmonic functions and corresponding coefficients. The series of scans which are averaged are defined for each subject session according to the closest 10 field map sessions in terms of date for scanner B whereas 10 acquisitions were acquired within a single session at scanner A which are averaged.

**Figure 1.**
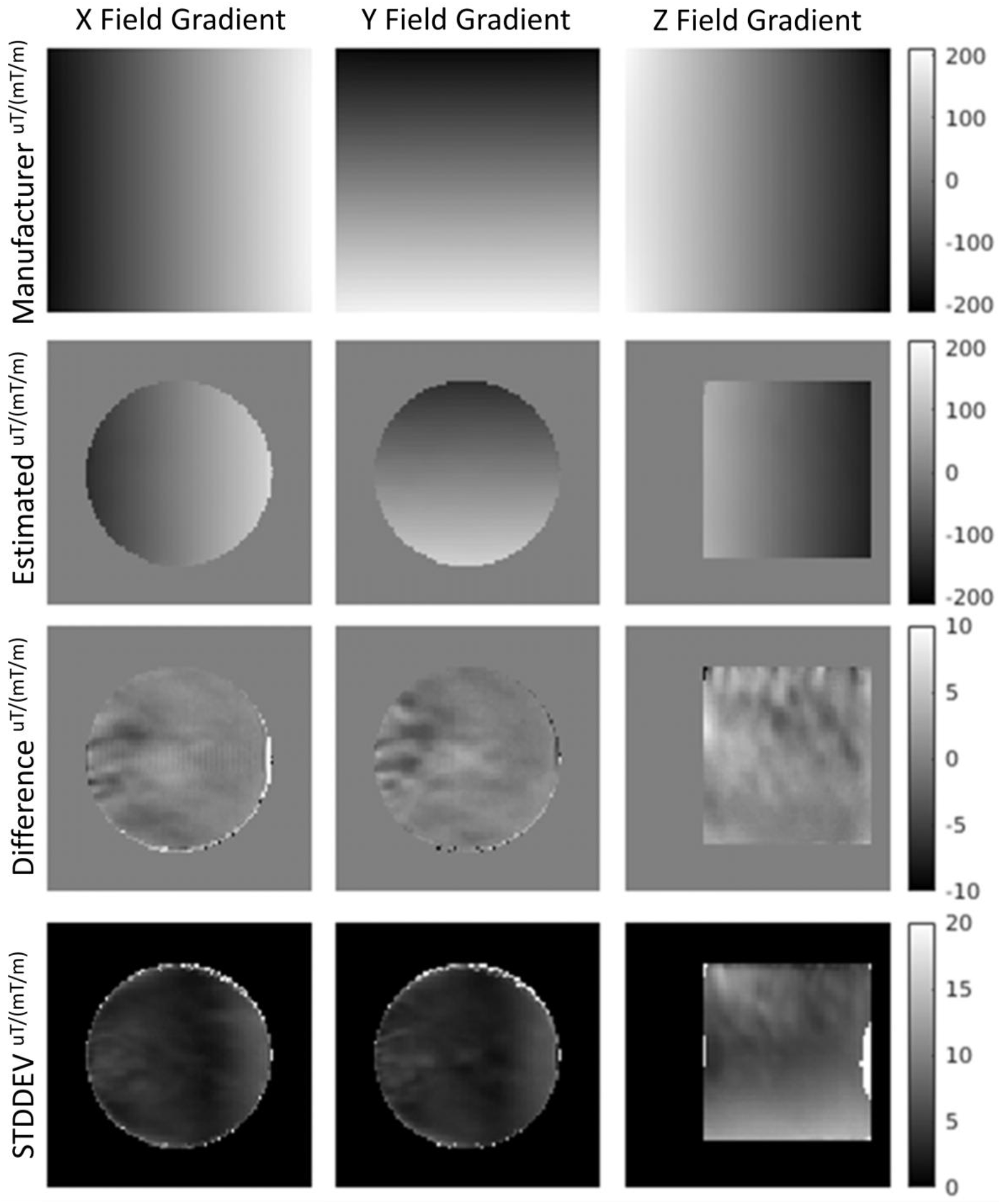
Here we show the manufacturer specified fields (top), the averaged empirically estimated (directly measured) fields (middle-top), the difference between these (middle-bottom), and the standard deviation in the empirically estimated fields across time (bottom) in units of uT (per mT/m of applied gradient). The field of view is 384mm by 384mm, and a mask is applied to the fields according the usable regions within the oil phantom (135mm radius from isocenter). The x and y magnetic field gradients are shown as an axial slices at isocenter (192mm), and the z magnetic field gradient is shown as a sagittal slice at isocenter (192mm).

### Estimating achieved b-values and gradient directions

A spatially varying tensor *L* relates the achieved magnetic field gradient to the intended one [12]:

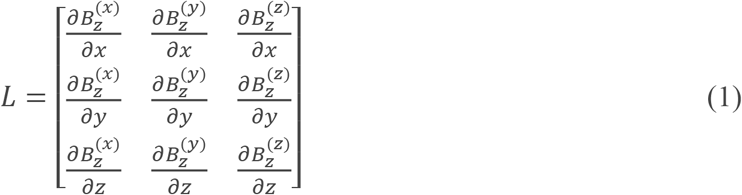

where 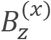 is the z component of the magnetic field produced by unit amplitude of a nominal x-gradient coil current, and similarly for (y) and (z). This tensor may be computed analytically from the solid harmonic approximation to the measured field, then evaluated at spatial locations of interest. We can use *L* to relate the assumed gradient vector to the achieved gradient field and as well as the assumed b-value to the achieved one. If we assume |*g*| = 1 then the adjusted gradient vector and b-value become:

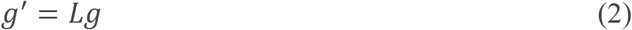

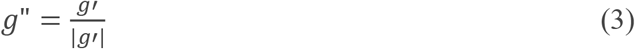

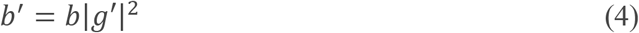

where *b*′ is the adjusted b-value and *g*” is the adjusted and normalized gradient vector. In the common situation where the scanner reports the intended gradient direction and amplitude but the full b-matrix [34–36] is not known, an approximate correction to adjust the signal *S*_*i*_ for the *i*^th^ diffusion acquisition relative to the reference signal *S*_0_ is [18]:

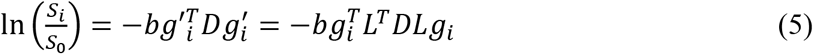

where *b* is the scalar b-value, *g* is the intended gradient vector, *g*′ is the actual gradient vector, and *D* is the diffusion tensor. If we substitute with *b*′ and *g*” equation 5 can be re-written as:

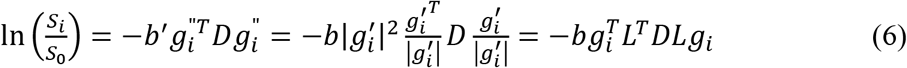

Importantly, this is spatially varying and processing occurs voxelwise, but this may be used in any desired way for further processing of the diffusion images. Figure 2 shows *L* for each voxel estimated using our empirical fieldmapping acquired on scanner B.

**Figure 2.**
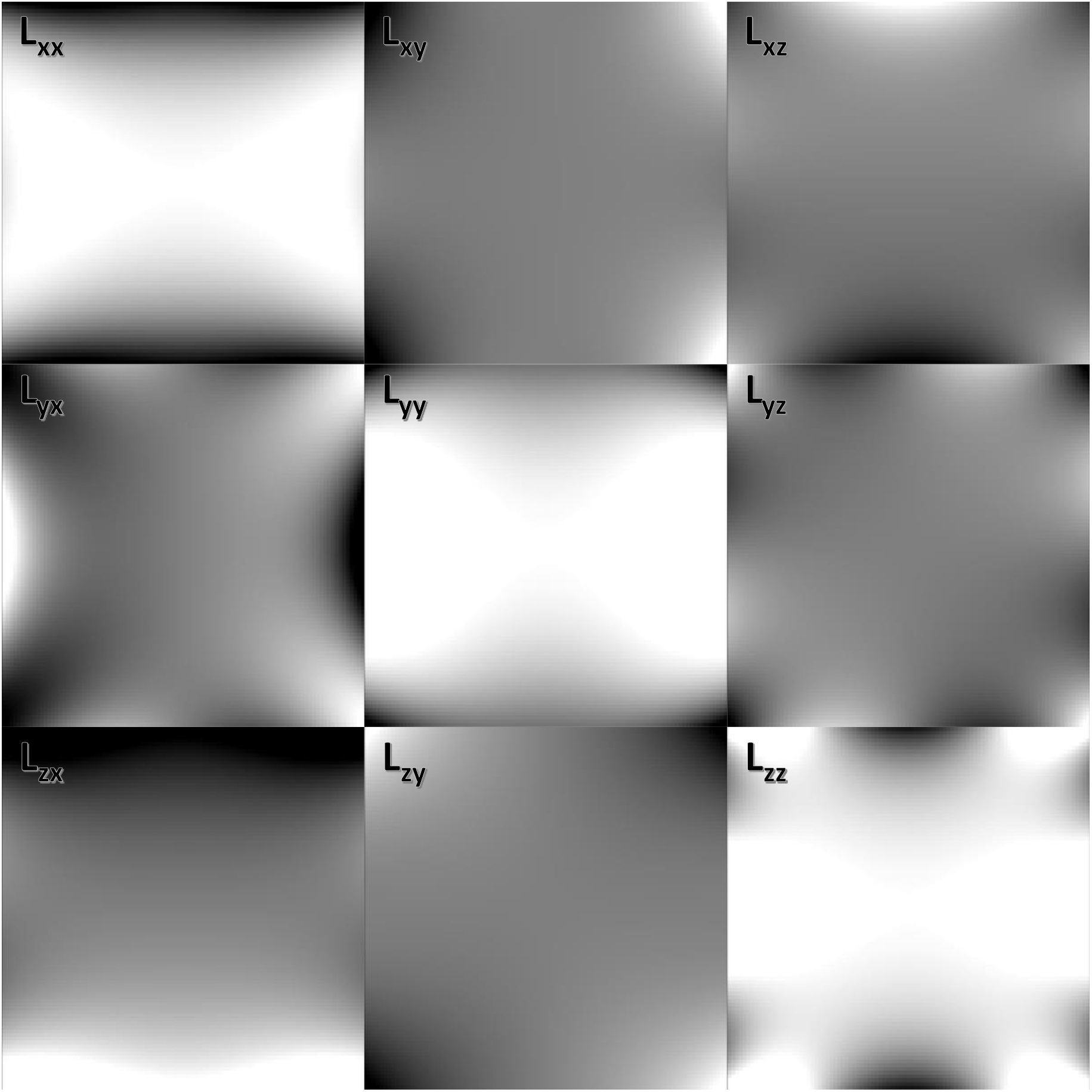
Gradient coil tensor L(r) (sagittal view) for each voxel position using 7^th^ order spherical harmonic expansion using only odd order terms. This was generated using the coefficients estimated from the empirical field mapping procedure.

## EXPERIMENTS

This section describes the set of analyses which aim to show the accuracy of the estimated fields as well as their impact on resulting DW-MRI metrics in phantom and human data. All DW-MRI are corrected for susceptibility distortion [37] and eddy current distortion [15] using FSL.

### Empirically Estimated Fieldmaps

Gradient nonlinearity correction is only viable if we can depend on the estimation to match the true fields. To investigate if the magnitude estimated fieldmaps closely approximate the true fields, we compare them to the fieldmaps specified by the manufacturer on scanner B. This was not done for scanner A as the manufacturer specifications for scanner A were not provided. For comparison, we take the average fieldmap from the latest 10 oil phantom scans on scanner B and calculate the voxel-wise difference between this and the manufacturer specified fields. To evaluate the stability of the empirical estimations, we report the variance across fields estimated from 40 individual oil phantom scans acquired over time on scanner B. These additional acquisitions are unnecessary for practical use and are strictly for evaluation purposes. Only a single acquisition would be needed for this method to be deployed on a scanner to be applied to all previous and future acquisitions. All evaluations on the empirical fields use a spherical mask with a radius of 135mm from isocenter.

### Polyvinylpyrrolidone (PVP) phantom

To evaluate the intra-scanner performance of the gradient field nonlinearity correction with the empirical fieldmaps in a controlled environment, we use a 43% Polyvinylpyrrolidone (PVP) aqueous solution in a sealed spherical container that is 160mm in diameter (PVP phantom) [38]. The PVP phantom is a large homogeneous material, and estimated metrics are expected to be the same across the entire volume. Additionally, toxicology has shown PVP to be safe for use, and PVP is stable and uniform. At scanner B, the phantom was scanned at three positions along the magnet axis: superior (4cm above isocenter), isocenter, and inferior (8cm below isocenter). At each position DWI data was acquired with diffusion weighting applied in twelve directions at a b-value of 1000 s/mm^2^ and twelve more were acquired at 2000 s/mm^2^ with a TR of 7775, a voxel resolution of 2.5mm by 2.5mm by 2.5mm, and a FOV of 240mm by 240mm by 170mm. Susceptibility distortion correction and eddy current distortion correction are applied without movement correction. Signal to noise ratio (SNR) was calculated by fitting the signal to a tensor in the phantom and taking the residuals after the fit. Using all diffusion volumes at each position, MD is calculated without and with gradient nonlinearity correction using the empirically derived fields and using the manufacturer specified fields. When calculating MD with the correction, the estimated achieved b-values and gradient directions for each voxel are used. We report error in terms of absolute percent error (APE) between each scan out of isocenter and the scan at isocenter. All non-diffusion volumes to a structural T1 image using a rigid body transform restricted to only use translations, and this registration is applied to the calculated MD before analysis.

### Human subject

To evaluate the intra-scanner and inter-scanner performance of the gradient field nonlinearity correction with the empirical fieldmaps in-vivo, we scanned a single subject at scanner A and scanner B. At scanner B, two sessions were acquired of the subject with one session acquired with the bridge of the subject’s nose positioned at isocenter within the magnet and one session acquired with the subject positioned 6cm superior from isocenter. At scanner A, only one session is acquired at isocenter. Each session consisted of twelve gradient directions at a b-value of 1000 s/mm^2^, twelve at a b-value of 2000 s/mm^2^, a TR of 3700ms, a voxel resolution of 2.5mm by 2.5mm by 2.5mm, and a FOV of 240mm by 240mm by 170mm. Susceptibility distortion correction and eddy current distortion correction are applied with movement correction for each session. Using all diffusion volumes from each session, MD is calculated without and with gradient nonlinearity correction using the empirically derived fields. At scanner B, MD is also calculated after correction with the manufacturer specifications. For analysis the scans are registered to a T1 acquired at isocenter using FSL Flirt [39]. We report MD error as the absolute percent error between the two scans acquired at scanner B and between the scan acquired at scanner A and the out of isocenter scan acquired at scanner B.

We also evaluate the performance of the empirical correction with higher quality acquisitions on scanner A. Again, two sessions are acquired of the subject: one with the bridge of the subject’s nose positioned at isocenter and one where the subject is shifted 4cm inferior from isocenter. Each session consisted of 384 gradient directions at a b-value of 1000 s/mm^2^, a voxel resolution of 2.5mm by 2.5mm by 2.5mm, and a FOV of 240mm by 240mm by 170mm. Susceptibility distortion correction and eddy current distortion correction are applied with movement correction for each session. Using all diffusion volumes from each session, MD is calculated without and with gradient nonlinearity correction using the empirically derived fields. For analysis the scans are registered to a T1 acquired at isocenter using FSL Flirt [39]. We report MD error as the absolute percent error between the two scans.

## RESULTS

### Empirically Estimated Fieldmaps

There are small differences between the manufacturer and the measured field produced by the gradient coil. These are shown in Figure 1 in units of uT scaled by the intensity (mT/m) of the applied gradient (uT/(mT/m), or mm). On average the difference at a given voxel is approximately 1 uT/(mT/m) in the x and y magnetic field gradients and 2 uT/(mT/m) in the z gradient field within 135mm of isocenter. The difference maps indicate the presence of some structural artifacts. The average standard deviation at a given voxel after 40 acquisitions acquired throughout a year is approximately 4 uT/(mT/m) in the x and y fields and 6 uT/(mT/m) in the z field within 135mm of isocenter.

### PVP phantom

The mean absolute percent error within the phantom between the inferior scan and the isocenter scan is approximately 5% before correction. After correction using the manufacturer fields, this falls to approximately 1.6%. Correcting with the empirically derived fields leads to 0.9% mean error. Figure 3 shows most of the error before correction in the inferior regions of the phantom which were furthest from isocenter in the inferior scan.

**Figure 3.**
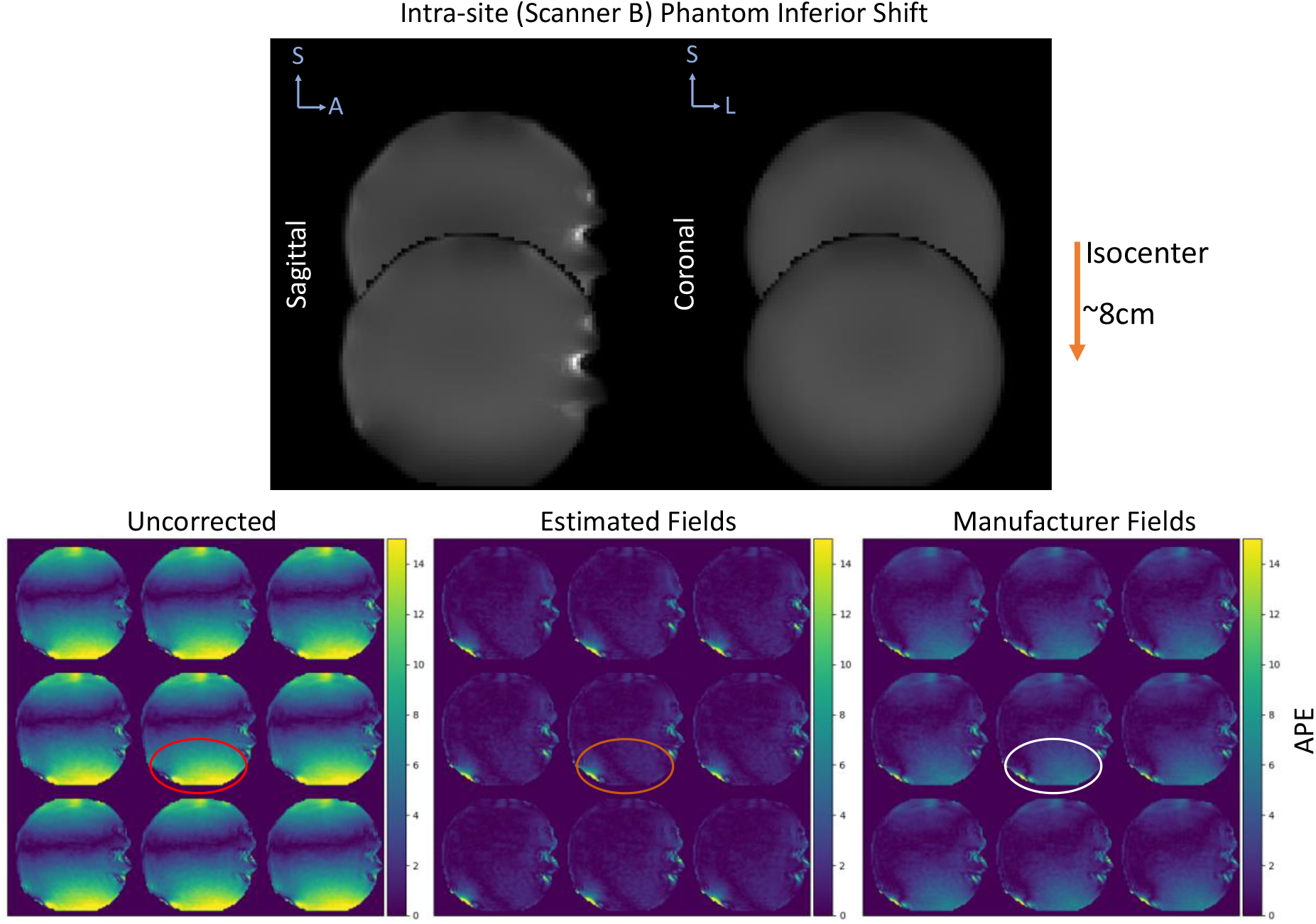
The absolute percent error (APE) in MD is shown for the PVP phantom with one session acquired at isocenter and another acquired 8cm inferior from isocenter. The top plot shows the sagittal and coronal view of the b0 from each session to demonstrate the shift within the scanner. The bottom plots show the APE for nine saggital slices before correction, after correction using the estimated fields, and after correction using the manufacturer specifications. The error before correction is most prominent in the inferior regions of the phantom as those were the furthest from isocenter during the second acquisition.

When uncorrected, the mean absolute percent error within the phantom between the superior scan and the isocenter scan is approximately 4.9%. After correction using the manufacturer fields, this falls to approximately 2%. Correcting with the empirically derived fields leads to 1.3% mean error. Figure 4 shows most of the error before correction in the superior regions of the phantom which were furthest from isocenter in the superior scan.

**Figure 4.**
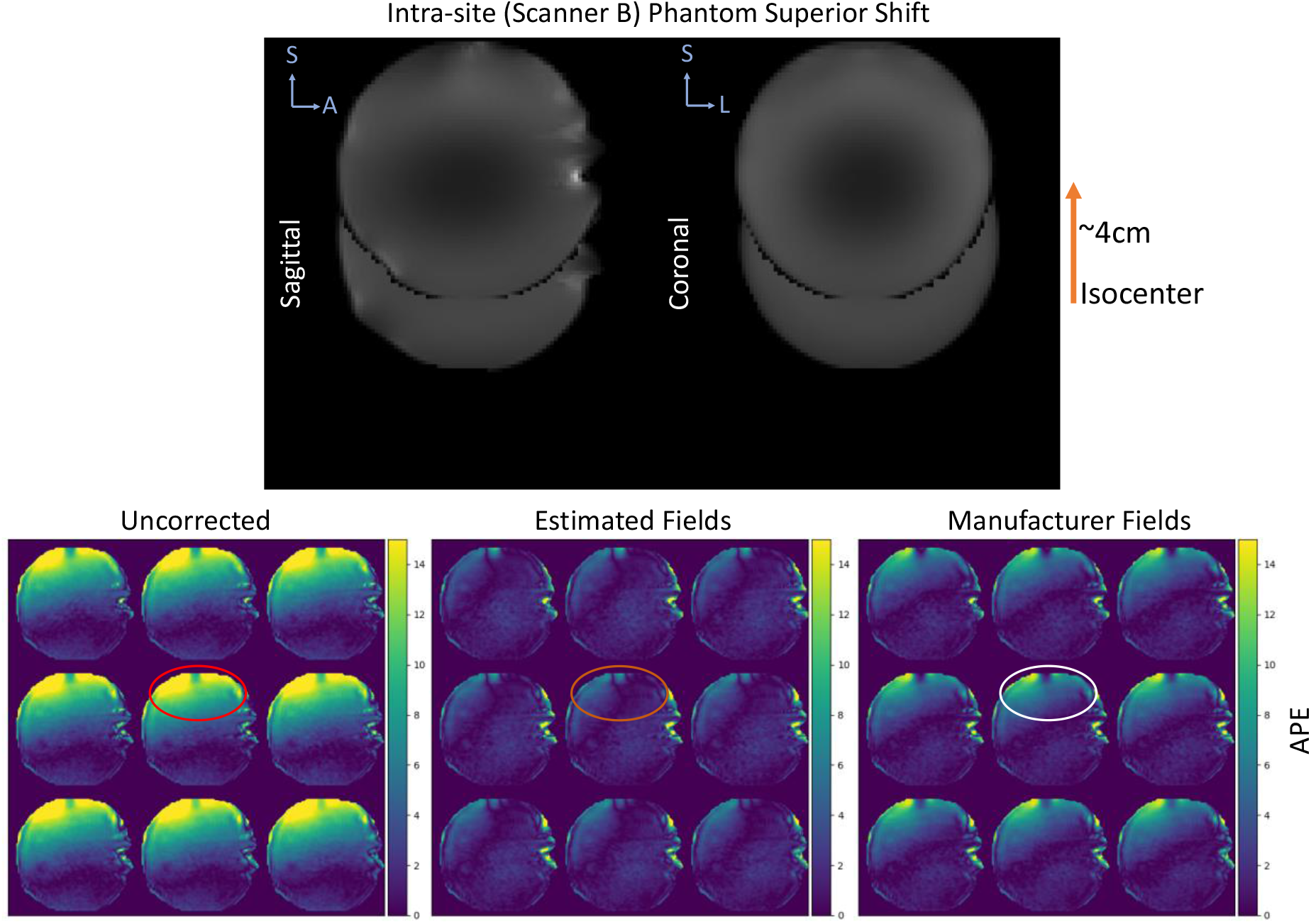
The absolute percent error (APE) in MD is shown for the PVP phantom with one session acquired at isocenter and another acquired 4cm superior from isocenter. The top plot shows the sagittal and coronal view of the b0 from each session to demonstrate the shift within the scanner. The bottom plots show the APE for nine saggital slices before correction, after correction using the estimated fields, and after correction using the manufacturer specifications. The error before correction is most prominent in the superior regions of the phantom as those were the furthest from isocenter during the second acquisition.

### Human repositioned

The intra-scanner sessions on scanner B result in a mean absolute percent error of 5.9% before correction within the brain volume excluding CSF regions. After correcting the scans using the empirically estimated fields, the mean error is reduced to 5.6% and further to 5.4% if the manufacturer specifications are used during correction. Just as in with the phantom, the error attributable to the gradient nonlinearities before correction appears in the superior regions of the brain which were furthest from isocenter during one of the sessions (Figure 5).

**Figure 5.**
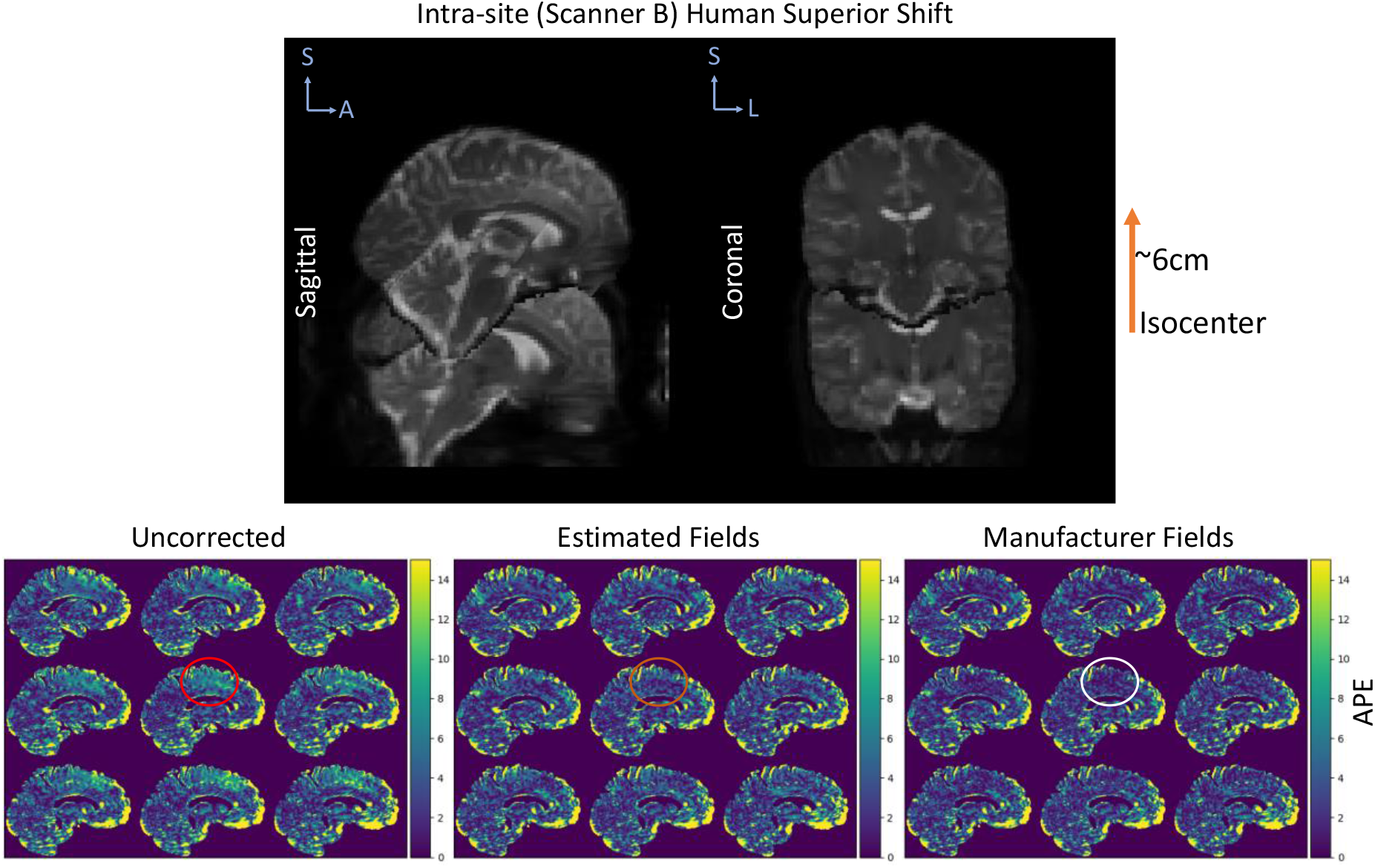
The absolute percent error (APE) in MD is shown for the human subject with one session acquired at isocenter and another acquired 6cm superior from isocenter on scanner B. The top plot shows the sagittal and coronal view of the b0 from each session to demonstrate the shift within the scanner. The bottom plots show the APE for nine saggital slices before correction, after correction using the estimated fields, and after correction using the manufacturer specifications. The error before correction is most prominent in the superior regions of the phantom as those were the furthest from isocenter during the second acquisition.

For the inter-scanner experiment, the mean absolute percent error before correction is 7.2% and is reduced 6.9% after correction using the estimated fields. Clearly the error that is accounted for in the correction is the superior regions of the brain which were furthest from isocenter during the session acquired on scanner B (Figure 6).

**Figure 6.**
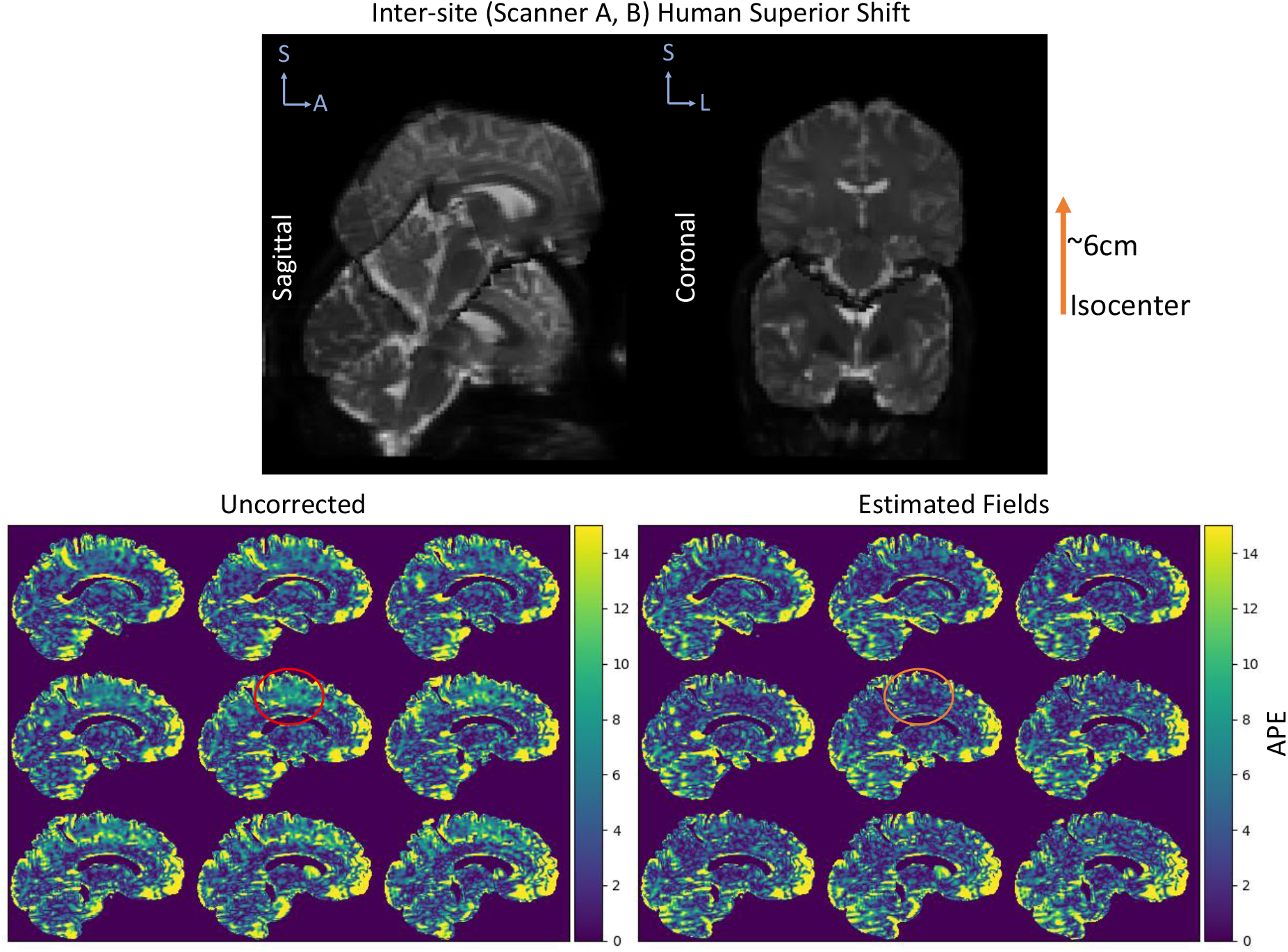
The absolute percent error (APE) in MD is shown for the human subject with one session acquired at isocenter on scanner A and another acquired 6cm superior from isocenter on scanner B. The top plot shows the sagittal and coronal view of the b0 from each session to demonstrate the shift within the scanner. The bottom plots show the APE for nine saggital slices before correction, after correction using the estimated fields, and after correction using the manufacturer specifications. The error before correction is most prominent in the superior regions of the phantom as those were the furthest from isocenter during the second acquisition.

The intra-scanner sessions acquired on scanner A using a significantly higher number of gradient directions results in a mean absolute percent error of 4.6% when no correction is applied. After correction using the empirically estimated fields, the mean error is reduced to 4.2%. The difference can be seen in the inferior regions of the brain, specifically the cerebellum which was furthest from isocenter during one of the sessions (Figure 7). Figure 8 shows the mean absolute percent error across all voxels within the phantom and within the brain volume excluding cerebrospinal fluid (CSF) regions for each method.

**Figure 7.**
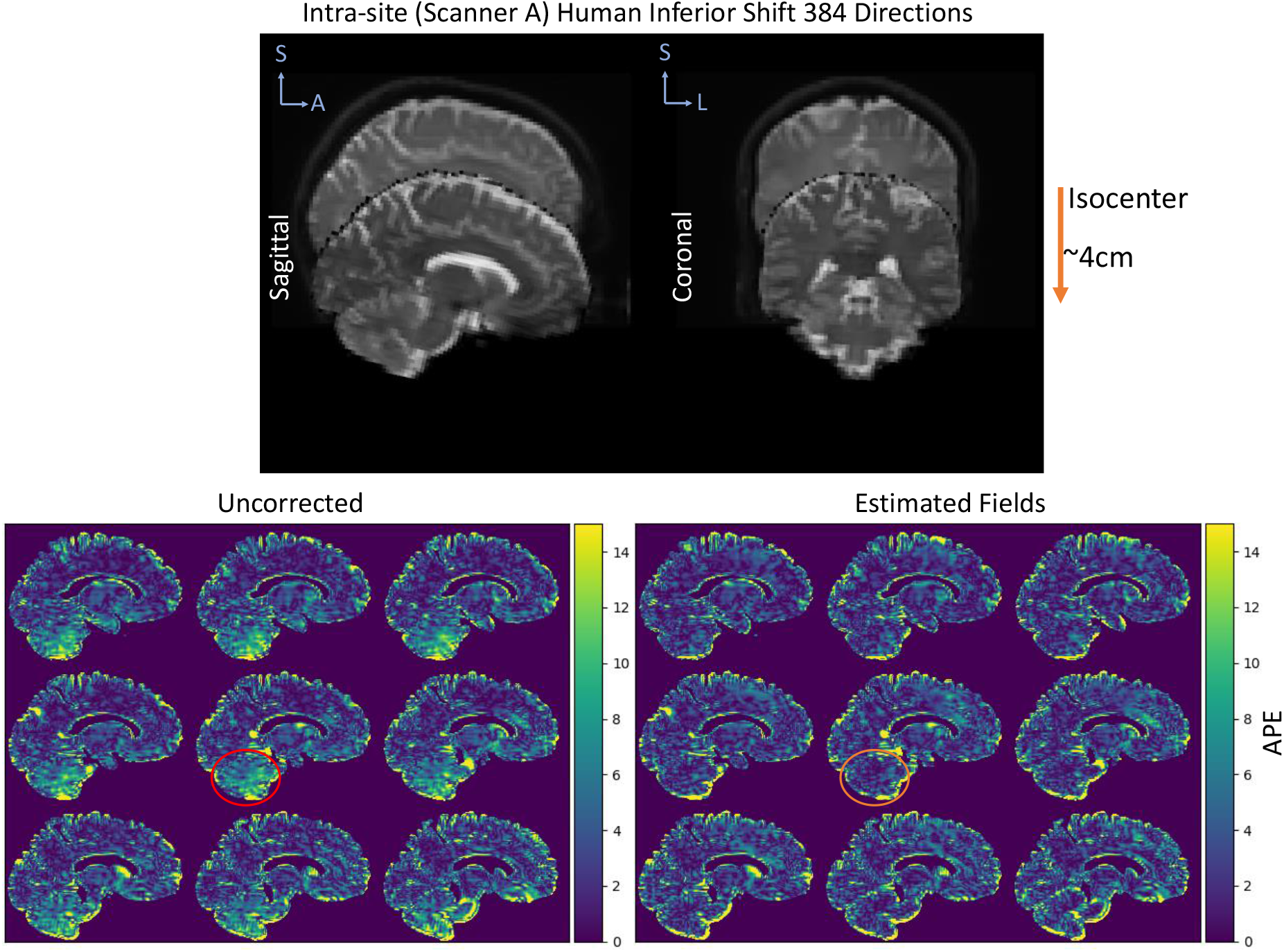
The absolute percent error (APE) in MD is shown for the human subject with one session acquired at isocenter and another acquired 4cm inferior from isocenter on scanner A. These acquisitions were acquired with 384 directions. The top plot shows the sagittal and coronal view of the b0 from each session to demonstrate the shift within the scanner. The bottom plots show the APE for nine saggital slices before correction, after correction using the estimated fields, and after correction using the manufacturer specifications. The error before correction is most prominent in the inferior regions of the phantom as those were the furthest from isocenter during the second acquisition.

**Figure 8.**
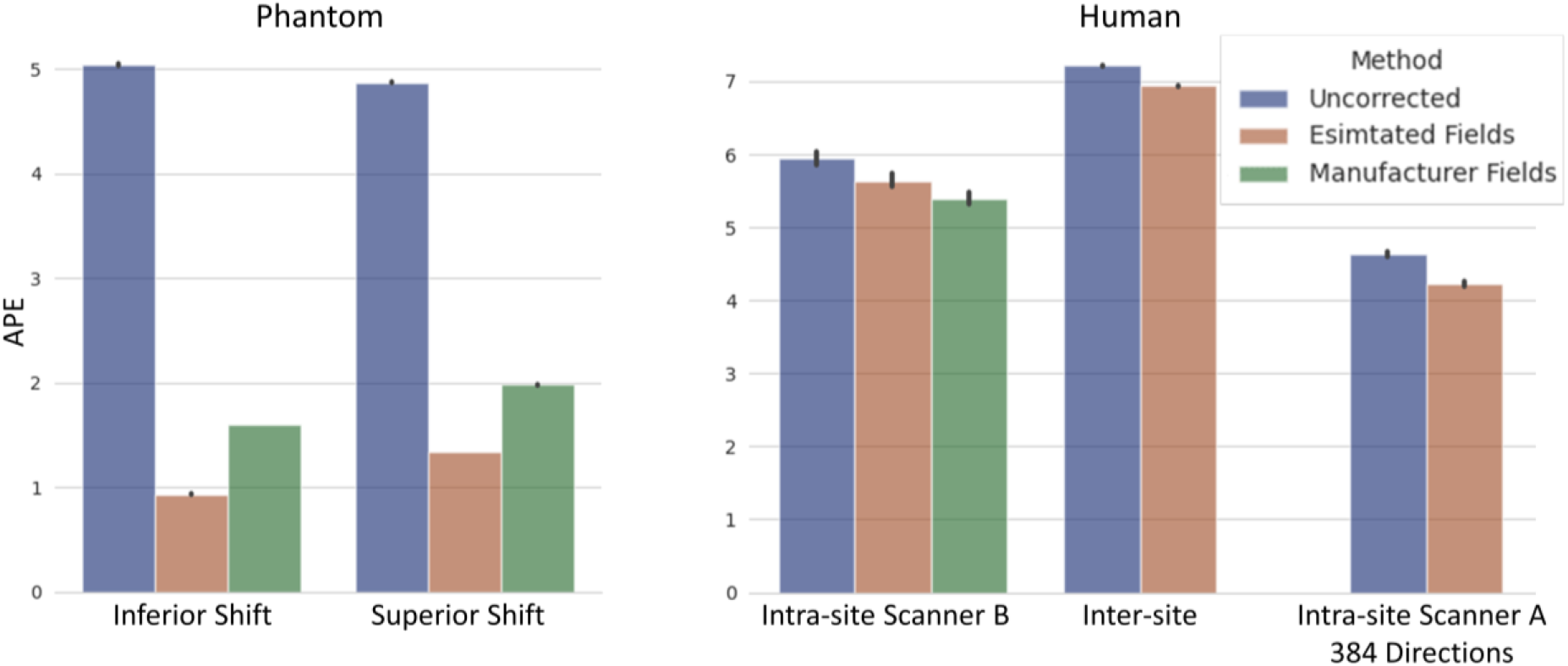
The mean APE within the phantom and brain excluding CSF regions are shown for each experiment without correction, after correction with the estimated fieldmaps, and after correction with the manufacturer specifications when available.

## DISCUSSION

In comparing the empirically estimated fields to the fields specified by the manufacturer, we find that our approximations are very similar. The largest differences are in the z gradient field which corresponds to the largest variations in all the estimated fields across 40 oil phantom acquisitions. In this study we use an average of fieldmaps across 10 acquisitions each acquired a week apart, but this should not be necessary as the field produced by the gradient coil depends only on the coil geometry and the current flowing in the coils. Unaltered system need only acquire the fields once for this method, but further study on the stability of the empirical mapping may be necessary. Additionally, further study on the stability of the fit of the spherical harmonics and the need for higher order basis may be necessary. Appendix A shows the effects of different orders.

The experiments with the PVP phantom show in a large isotropic volume the impact of the gradient nonlinearities within the magnet and the effectiveness of the correction. The small superior shift of 4cm results in over 15% error in the superior voxels. In the case of a large inferior shift and a smaller superior shift, the mean error is increased by a factor of two to five if these effects are not accounted for. If we consider the experiments involving the human subject, we can see the impact of this correction is reduced. This could in part due to imperfect registration which seems to have contributed to error in the anterior regions of the brain. Results may vary depending on registration strategy. We have tried multiple techniques with similar results. Though the absolute percent error only changed by 0.3% to 0.4%, some small regions see a similar magnitude of improvement, and it is qualitatively clear that the correction is impacting regions we expect. The differences between resulting absolute percent error using the empirical fields and the manufacturer fields is varies between the phantom and the human subject. The results for the phantom indicate that the estimated fields improve performance of the method, but the human subject results show a small advantage for using the manufacturer field directly.

Though all intra-scanner results on scanner B are compared against using the manufacturer field directly, future work should investigate the sensitivity of our proposed method and compare with other field mapping methods such as proposed by Janke et. al [24] even though these methods require that the manufacturer provide the solid harmonic coefficients. In recent work, another approach is proposed for correcting voxel-wise b-value errors. Instead of correcting for gradient nonlinearities in the coil, this method directly estimates a voxel-wise b-value map that is used to correct resulting diffusion metrics [40]. While this method could account for errors that stem from other sources of deviation than just gradient nonlinearities, the model requires an estimation of more parameters and likely it would be best practice to acquire a calibration scan along with every subject acquisition. In comparison to apply the approach proposed in this work, only a single calibration scan is necessary for each system.

While this method is successful in circumventing the need for manufacturer specifications which are not always readily available, it should be noted that vendor-provided on-scanner gradient nonlinearity correction is preferred for translation in a clinical environment. Additionally, when working with any DICOM data coordinating world coordinate frame and patient frames can be incredibly nuanced and should be considered carefully when applying any corrections post acquisition. However, our approach remains as a solution to correct retroactively to enable the use of acquired datasets which should be corrected for gradient nonlinearity effects for use in clinic and in research.

## CONCLUSION

This work shows that the errors caused by gradient nonlinearities is apparent in metrics derived from DW-MRI but can be reduced using the correction outlined by Bammer et al. Using empirically derived fields, we can achieve similar results without needing manufacturer specification of the hardware. In both phantom and in-vivo data, error in MD can be significantly reduced by applying this correction. We advocate for the use of gradient nonlinearity correction in standard diffusion preprocessing pipelines and provide a simple method for empirically measuring the fields necessary to account for the achieved b-values and b-vectors.

## ACKNOWLEDGEMENTS

This work was supported by the National Institutes of Health under award numbers R01EB017230 and T32EB001628, and in part by the National Center for Research Resources, Grant UL1 RR024975-01. The content is solely the responsibility of the authors and does not necessarily represent the official views of the NIH.

## APPENDIX A

The PVP phantom is corrected using fieldmaps estimated with various orders of solid harmonics. Regardless of the order, both FA and MD reproducibility errors decrease when compared to the uncorrected error. However, we find that a 3^rd^ order basis results in the lowest FA error but a higher MD error. Between the higher order basis, the 7^th^ order solid harmonics achieves lower FA error.

**Figure A.1:**
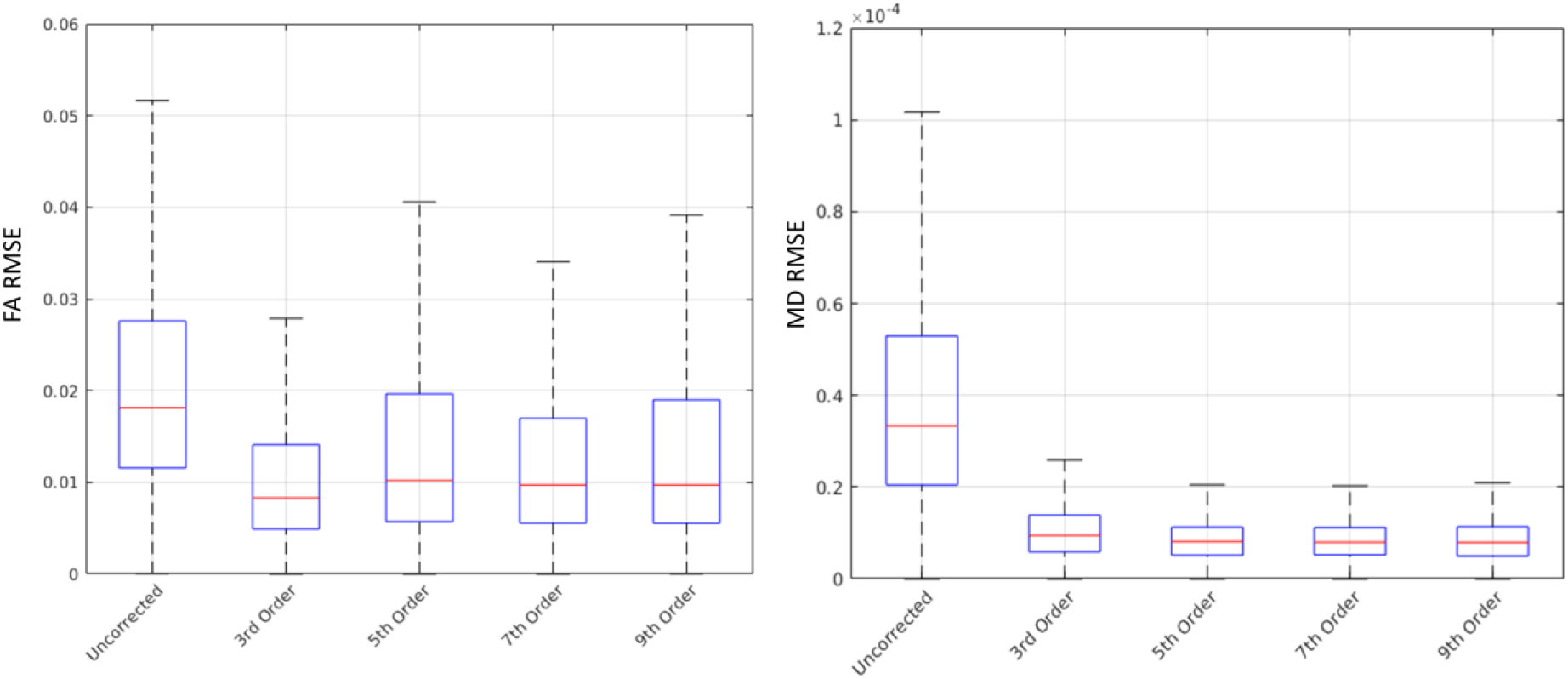
The reproducibility error in FA and MD for the PVP phantom are calculated using the estimated fieldmap utilizing different orders of solid harmonics. Orders higher than 3^rd^ achieve lower MD RMSE but tend to have higher FA RMSE.

## REFERENCES

1. Glover, G.H. and N.J. Pelc, Method for correcting image distortion due to gradient nonuniformity. 1986, Google Patents.

2. Michiels, J., et al., On the problem of geometric distortion in magnetic resonance images for stereotactic neurosurgery. Magnetic resonance imaging, 1994. 12(5): p. 749–765.

3. Sumanaweera, T., et al., Quantifying MRI geometric distortion in tissue. Magnetic resonance in medicine, 1994. 31(1): p. 40–47.

4. Langlois, S., et al., MRI geometric distortion: a simple approach to correcting the effects of non-linear gradient fields. Journal of Magnetic Resonance Imaging: An Official Journal of the International Society for Magnetic Resonance in Medicine, 1999. 9(6): p. 821–831.

5. LeBihan, D. and R. Tumer, Diffusion and perfusion, magnetic resonance imaging, Mozbey Year Book. 1992, Inc.

6. Conturo, T.E., et al., Diffusion MRI: precision, accuracy and flow effects. NMR in Biomedicine, 1995. 8(7): p. 307–332.

7. Bernstein, M.A. and J.A. Polzin, Method and system for correcting errors in MR images due to regions of gradient non-uniformity for parametric imaging such as quantitative flow analysis. 2000, Google Patents.

8. Bammer, R., et al., Assessment of spatial gradient field distortion in diffusion-weighted imaging. Proceedings of the International Society for Magnetic Resonance in Medicine , Honolulu, HI, 2002: p. 1172.

9. Robson, M. Non-linear gradients on clinical MRI systems introduce systematic errors in ADC and DTI measurements. in Proceedings of the 10th Annual Meeting of ISMRM, Honolulu. 2002.

10. Basser, P.J., Inferring microstructural features and the physiological state of tissues from diffusion-weighted images. NMR in Biomedicine, 1995. 8(7): p. 333–344.

11. Frank, L.R., Anisotropy in high angular resolution diffusion-weighted MRI. Magnetic Resonance in Medicine: An Official Journal of the International Society for Magnetic Resonance in Medicine, 2001. 45(6): p. 935–939.

12. Bammer, R., et al., Analysis and generalized correction of the effect of spatial gradient field distortions in diffusion-weighted imaging. Magnetic Resonance in Medicine: An Official Journal of the International Society for Magnetic Resonance in Medicine, 2003. 50(3): p. 560–569.

13. Setsompop, K., et al., Pushing the limits of in vivo diffusion MRI for the Human Connectome Project. Neuroimage, 2013. 80: p. 220–233.

14. Malyarenko, D.I., et al., Demonstration of nonlinearity bias in the measurement of the apparent diffusion coefficient in multicenter trials. J Magnetic resonance in medicine, 2016. 75(3): p. 1312–1323.

15. Andersson, J.L. and S.N. Sotiropoulos, An integrated approach to correction for off-resonance effects and subject movement in diffusion MR imaging. Neuroimage, 2016. 125: p. 1063–1078.

16. McNab, J.A., et al., The Human Connectome Project and beyond: initial applications of 300 mT/m gradients. Neuroimage, 2013. 80: p. 234–245.

17. Mesri, H.Y., et al., The adverse effect of gradient nonlinearities on diffusion MRI: From voxels to group studies. NeuroImage, 2019: p. 116127.

18. Tan, E.T., et al., Improved correction for gradient nonlinearity effects in diffusion-weighted imaging. Journal of Magnetic Resonance Imaging, 2013. 38(2): p. 448–453.

19. Newitt, D.C., et al., Gradient nonlinearity correction to improve apparent diffusion coefficient accuracy and standardization in the american college of radiology imaging network 6698 breast cancer trial. Journal of Magnetic Resonance Imaging, 2015. 42(4): p. 908–919.

20. Malyarenko, D.I., B.D. Ross, and T.L. Chenevert, Analysis and correction of gradient nonlinearity bias in apparent diffusion coefficient measurements. Magnetic resonance in medicine, 2014. 71(3): p. 1312–1323.

21. Malyarenko, D.I. and T.L. Chenevert, Practical estimate of gradient nonlinearity for implementation of apparent diffusion coefficient bias correction. Journal of Magnetic Resonance Imaging, 2014. 40(6): p. 1487–1495.

22. Sotiropoulos, S.N., et al., Advances in diffusion MRI acquisition and processing in the Human Connectome Project. Neuroimage, 2013. 80: p. 125–143.

23. Glasser, M.F., et al., The minimal preprocessing pipelines for the Human Connectome Project. Neuroimage, 2013. 80: p. 105–124.

24. Janke, A., et al., Use of spherical harmonic deconvolution methods to compensate for nonlinear gradient effects on MRI images. Magnetic Resonance in Medicine: An Official Journal of the International Society for Magnetic Resonance in Medicine, 2004. 52(1): p. 115–122.

25. Doran, S.J., et al., A complete distortion correction for MR images: I. Gradient warp correction. Physics in Medicine & Biology, 2005. 50(7): p. 1343.

26. Tao, S., et al., Integrated image reconstruction and gradient nonlinearity correction. Magnetic resonance in medicine, 2015. 74(4): p. 1019–1031.

27. Tao, A.T., et al., Improving apparent diffusion coefficient accuracy on a compact 3T MRI scanner using gradient nonlinearity correction. Journal of Magnetic Resonance Imaging, 2018. 48(6): p. 1498–1507.

28. Tao, S., et al., NonCartesian MR image reconstruction with integrated gradient nonlinearity correction. Medical physics, 2015. 42(12): p. 7190–7201.

29. Rogers, B.P., et al. Phantom-based field maps for gradient nonlinearity correction in diffusion imaging. in Medical Imaging 2018: Physics of Medical Imaging. 2018. International Society for Optics and Photonics.

30. Rogers, B.P., et al. Stability of gradient field corrections for quantitative diffusion MRI. in Medical Imaging 2017: Physics of Medical Imaging. 2017. International Society for Optics and Photonics.

31. Tough, R.J. and A.J. Stone, Properties of the regular and irregular solid harmonics. Journal Of Physics A: Mathematical General, 1977. 10(8): p. 1261.

32. Caola, M., Solid harmonics and their addition theorems. Journal of Physics A: Mathematical General, 1978. 11(2): p. L23.

33. Makris, N., et al., MRI-based anatomical model of the human head for specific absorption rate mapping. Medical & biological engineering & computing, 2008. 46(12): p. 1239–1251.

34. Mattiello, J., P.J. Basser, and D. Le Bihan, The b matrix in diffusion tensor echo-planar imaging. Magnetic Resonance in Medicine, 1997. 37(2): p. 292–300.

35. Mattiello, J., P.J. Basser, and D. LeBihan, Analytical expressions for the b matrix in NMR diffusion imaging and spectroscopy. Journal of magnetic resonance, Series A, 1994. 108(2): p. 131–141.

36. Alger, J.R., The diffusion tensor imaging toolbox. Journal of Neuroscience, 2012. 32(22): p. 7418–7428.

37. Andersson, J.L., S. Skare, and J. Ashburner, How to correct susceptibility distortions in spin-echo echo-planar images: application to diffusion tensor imaging. Neuroimage, 2003. 20(2): p. 870–888.

38. Pierpaoli, C., et al. Polyvinylpyrrolidone (PVP) water solutions as isotropic phantoms for diffusion MRI studies. in Proc Intl Soc Magn Reson Med. 2009.

39. Jenkinson, M., et al., Improved optimization for the robust and accurate linear registration and motion correction of brain images. Neuroimage, 2002. 17(2): p. 825–841.

40. Lee, Y., et al., A comprehensive approach for correcting voxel-wise b-value errors in diffusion MRI. 2019.

